# CARPDM: cost-effective antibiotic resistome profiling of metagenomic samples using targeted enrichment

**DOI:** 10.1101/2024.03.27.587061

**Authors:** Dirk Hackenberger, Hamna Imtiaz, Amogelang R. Raphenya, Brian P. Alcock, Hendrik N. Poinar, Gerard D. Wright, Andrew G. McArthur

## Abstract

Better interrogation of antimicrobial resistance requires new approaches to detect the associated genes in metagenomic samples. Targeted enrichment is an ideal method for their sequencing and characterization. However, no open-source, up-to-date hybridization probe set targeting antimicrobial resistance genes exists. Here we describe CARPDM, a probe design software package made to run alongside all future Comprehensive Antibiotic Resistance Database releases. To test its efficacy, we have created and validated two separate probe sets: AllCARD, which enriches all genes encoded in the Comprehensive Antibiotic Resistance Database’s protein homolog models (n = 4,661), and clinicalCARD, which focuses on a clinically relevant subset of resistance genes (n = 323). We demonstrate that allCARD increases the number of reads mapping to resistance genes by up to 594-fold. ClinicalCARD performs similarly when clinically relevant genes are present, increasing the number of resistance-gene mapping reads by up to 598-fold. In parallel with this development, we have established a protocol to synthesize any probe set in-house, saving up to 350 dollars per reaction. Together, these probe sets, their associated design program CARPDM, and the protocol for in-house synthesis will democratize metagenomic resistome analyses, allowing researchers access to a cost-effective and efficient means to explore the antibiotic resistome.

## Introduction

Antimicrobial resistance (AMR) is a growing and global problem. In 2019, AMR was estimated to be directly responsible for 1.27M deaths (1). By 2050, this number may be as high as 10M (2). Most of this impact is and will continue to be in regions least equipped to combat it, largely due to a lack of resources (1). Therefore, it is imperative that we devise cost-effective solutions to address AMR more effectively and equitably. Bacterial evolution has been marked by the arms race of antibiotics and their associated resistance genes, providing a competitive edge to their producers (3–8). This struggle is revealed in the vast reservoir of antimicrobial resistance genes (ARGs) in environmental microbes, requiring only mobilization through horizontal gene transfer to be effective against any treatment we deploy (9–12). Therefore, to better combat AMR emergence, we need to improve how we detect the full complement of ARGs (the resistome(13)) in environmental and other reservoirs. Attractive monitoring targets are environments such as wastewater, a fertile ground for genetic exchange between bacteria [5] that provides a snapshot of ARG prevalence in the community (14–20). Natural environments such as soils, rivers, and farms are known reservoirs of ARGs, many with the potential to mobilize into pathogens (21–25). Finally, profiling the resistome of human and animal microbiomes allows us to identify critical determinants in the spread of resistance within and between these two groups (26, 27).

Investigating these rich data sources requires methods to characterize their resistome. Several techniques exist that may fill this niche, each with its limitations. PCR, for example, is commonly used to detect ARGs in *Mycobacterium tuberculosis* isolates (28). While useful for a small, targeted set of genes, the Comprehensive Antibiotic Resistance Database (CARD) (29) hosts over 5000 resistance determinants, an untenable number for PCR methods. Furthermore, because of the specificity of PCR, there is little chance of detecting distantly related genes, as a single nucleotide substitution may eliminate any signal from the assay. Finally, even if one detects a novel sequence variant by PCR, without follow-up amplicon sequencing, there is no way to identify it. This makes phylogenetic tracking of the spread of AMR far more difficult.

Shotgun DNA sequencing can detect all the genes in a sample given sufficient depth. Groups have used this method to characterize environmental (25, 30) and worldwide wastewater (14, 15) resistomes. A limitation of this technique is that all ARGs in a metagenomic sample typically represent <1% of the total DNA. Individual ARGs may be several orders of magnitude less than that in abundance. For example, a single 1kb ARG that makes up 1x10^-6^% of the DNA in a sample would require 10 Gbp of sequence data to obtain 10-fold coverage of the ARG. Performing this work on a NextSeq 2000 would cost over USD$1500. As such, while deep sequencing can be used to characterize metagenomic resistomes, it entails a high cost per sample, most of which will be spent sequencing background DNA. The associated volume of data also increases equipment costs and computing power needed to parse the data, further constraining this technique’s use in resource-limited settings (31).

Targeted enrichment is a modification to shotgun sequencing that allows robust detection of a broad range of specific, low-abundance targets with less sequencing. In this protocol, DNA from a sequencing library is denatured, allowing biotinylated RNA ‘probes’ complementary to a set of target sequences to hybridize (32, 33). Streptavidin-coated magnetic beads capture these biotinylated RNA probes and their complementary DNA partners from the background (Figure 1). This process increases the proportion of the target DNA in a library, allowing one to sequence less yet detect more.

**Figure 1:**
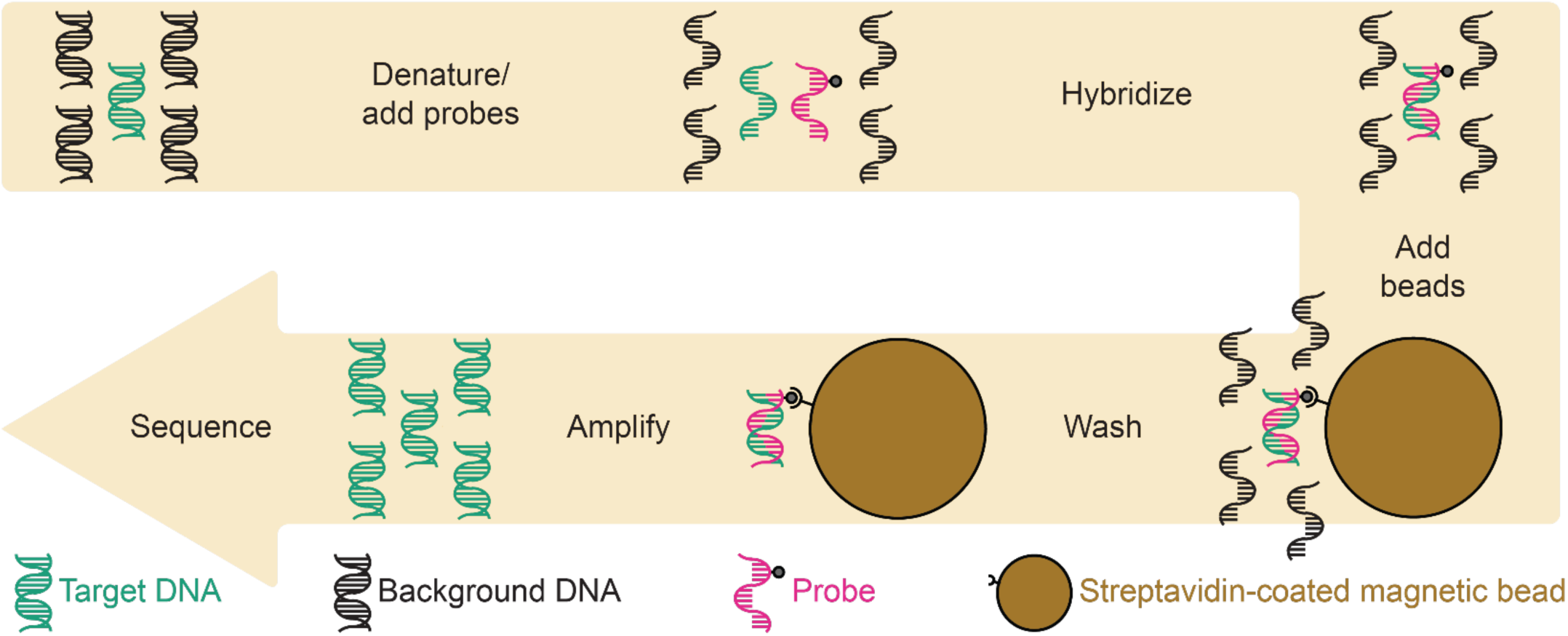
Targeted enrichment workflow. Magnetic streptavidin beads are used to bind biotinylated RNA molecules, which are, in turn, attached to a complementary DNA partner. One can considerably bias the resulting library towards a target fraction by pelleting the beads and washing away the background before amplification and sequencing.

A probe-capture protocol is ideal for detecting thousands of ARGs with a fraction of the sequencing required by brute-force shotgun approaches. However, two challenges exist for the probe-capture strategy in the AMR space. First, there is no up-to-date and open-source AMR probe set, as the most recent was designed against only 2,021 ARGs from CARDv1.0.1, released in 2015 (34). The second is that the cost of probes from commercial suppliers can be up to $350 per reaction, diminishing cost savings relative to shotgun sequencing. To address the first of these challenges, we have written a software package – the Comprehensive Antibiotic Resistance Probe Design Machine (CARPDM) - to generate a stringently filtered probe set with minimal off-target enrichment from the CARD v3.2.5 protein homolog model (allCARD) ARGs. This software package will run alongside all new releases of CARD, ensuring there is always an open-source and up-to-date probe set to enrich ARGs from any sample. We also curated a list of 323 clinically relevant ARGs and generated a smaller probe set (clinicalCARD) with the same program, providing a more focused alternative to the allCARD probe set, as the full complement of ARGs may not be necessary for many projects in healthcare settings. To address the cost of commercial probes, we have developed a protocol that allows in-house synthesis of any probe set from a Twist Biosciences^®^ oligo-pool. With this strategy, researchers can synthesize thousands of reactions worth of any probe set for a one-time fee lower than that of a typical 16-reaction kit from a commercial supplier.

For validation of our probe sets and in-house synthesis of probes, five conditions were tested on two wastewater and three soil samples in duplicate, comparing enrichment via the 2015 CARD v1.0.1 (34), allCARD, clinicalCARD probe sets to sequencing without enrichment, plus examination of the comparative performance of commercially supplied probes and probes synthesized using our new protocol. We demonstrate that our in-house synthesized probe set detected more ARGs with the same amount of sequencing on the samples we tested when compared to a commonly used commercial option. We also illustrate CARPDM’s capability as a probe design platform by validating allCARD and clinicalCARD probe sets on wastewater and soil samples. We found that allCARD detects far more ARGs than the probe set generated against CARD v1.0.1 (34). Finally, we show that clinicalCARD detects clinically relevant ARGs with less sequencing and better coverage than allCARD.

## Methods

### ARG Selection for AllCARD

Only ARGs curated as protein homolog models were included from CARD v3.2.5 (n = 4,661), as these ARGs do not confer resistance via the acquisition of mutations (i.e., CARD’s protein variant models). Including mutation-based ARGs in the design would enrich wild-type alleles, as enrichment probes can readily hybridize over a single mutation, diluting sequencing effort.

### ARG Selection for ClinicalCARD

The CARD prevalence data v4.0.0 [19] aided in identifying clinically relevant ARGs. This dataset is a collection of 221,175 sequencing assemblies from 377 pathogens with associated ARGs identified by CARD’s Resistome Gene Identifier RGI software (35). These data include a collection of 21,079 completely sequenced chromosomes and 41,828 completely sequenced plasmids. Preliminary examination of these data illustrated that the distribution of ARGs was almost binary, i.e., each ARG predominantly occurred in either plasmids or chromosomes, but not both (Supplemental Figure 1). It also showed that ARGs with a higher occurrence on plasmids were far more likely to be clinically relevant (36–39). As such, the first list of candidate clinically relevant ARGs included any ARG in CARD with at least ten occurrences in plasmids, yielding 237 ARGs. Any ARG with >5% but <95% prevalence in any ESKAPE pathogen (40, 41) was also added to this list, yielding a further 124 ARGs. Finally, 42 ARGs identified as clinically significant via a literature review were added (42, 43), resulting in a draft set of 403 total ARGs. This set was manually curated based on CARD prevalence data, considering the species they appeared in, the genomic context (whether it was mobile, i.e., on a plasmid), the drug classes they impact, the environment of the host organism, and the relative risk each species imposed as a pathogen. Our approach was developed following recent ARG risk frameworks developed for metagenomic data (42, 43). In short, our manual curation included mobile ARGs that confer resistance to a clinical antibiotic, are present in a human pathogen, and share an environment with humans or animals. After curation, 323 ARGs were deemed clinically relevant and included in the clinicalCARD probe set design.

### Probe Design

CARPDM is written in Python, save for the first step, which employs the program BaitsTools (44) to tile probe sequences along ARG sequences. Probes designed by CARPDM have an 80 nt length as per the prior validated ARG bait set for CARD v1.0.1. This is also an ideal size for adding amplification primers while remaining below the 120nt cut-off for the base level of a Twist oligo-pool if synthesizing probes in-house. The probes also have an extremely high density before filtering, with a tiling distance of 4 nt along ARG nucleotide sequences. This increases redundancy in the probe set and makes it more robust against stringent downstream filtering. As input, the allCARD design used all 4,661 CARD v3.2.5 protein homolog model ARGs, while clinicalCARD’s design used the curated 323 clinically relevant ARGs.

There are three filtering steps after the initial probe construction (Supplemental Figure 2). The first is a filter based on sequence, in which the probes are deduplicated, confirmed as 80 nt, contain no ambiguous bases, have no perfect complements within the set, have a Tm >50°C, and contain no LguI restriction endonuclease cut sites. The last condition is essential for the synthesis protocol. Since bacteria are likely to be the most abundant organism in most metagenomic samples (45), the second filter is a BLASTN (46) search against the nt database to minimize off-target enrichment and removes probes with >80% identity over <50 nt to any bacterial sequence, as any probe similar to a bacterial sequence over >50 nt is likely the ARG itself. This filter also removes probes with >80% similarity over >50 nt to any viral, archaeal, or eukaryotic sequence unless they match perfectly (i.e., likely sequence contamination by an ARG).

The last filter removes redundancy in the probe set, using BLASTN to compare the probes against each other. To remove probes that could bind to each other rather than the target DNA, any probes found to be complementary over the entire 80 nt are collapsed to a single representative. After removing complementary probes, CARPDM determines if the number of probes in the set is below a pre-defined cut-off. For allCARD, we arbitrarily set this cut-off at 40,000 probes, while for clinicalCARD, it was set to 20,000 probes. If not below the cut-off, CARPDM collapses probes with 79 nt of complementarity to a single representative, repeating the process with decreasing length of complementarity until below the cut-off.

### *In Silico* Analysis of Probe Sets

BLASTN (46) was used to compare the probe sets to the ARG sequences. Using Python and NumPy (47), “target” arrays of zeros were constructed. Each of these arrays represented a single ARG against which the probeset was designed. After aligning all probes against the input set of target sequences with BLASTN, arrays of ones (matches) and zeros (mismatches) were constructed for each probe:target alignment. These arrays were then added to the corresponding target array at the start position of the alignment. Repeating this for every probe:target alignment yields the tallied coverage of each nucleotide position by probes. This coverage array was used to compute summary statistics.

### Probe Synthesis

For in-house synthesis of probes, the first step is to amplify an oligo pool by PCR. For this to happen, two PCR primers must flank each probe sequence, with one including the T7 transcription start site for RNA synthesis and the other including an LguI restriction endonuclease digest sequence so that it can be cleaved after amplification (Supplemental Figure 3). In support of in-house synthesis, CARPDM first appends a T7 transcription start site with three extra guanines to one end of every probe sequence to reduce transcription efficiency variability (48) and then creates every possible primer with a terminal LguI cut site of the correct length to yield a final oligo of 120 nt when appended to the opposite side. It compares these putative primers to the concatenated T7/probe sequences using BLASTN and selects the primer with the fewest matches as the second amplification primer, appending the reverse complement to the opposite side of each probe sequence. The resulting sequences were ordered as an oligo-pool from Twist Biosciences (San Francisco, CA), and the amplification primers were ordered from Integrated DNA Technologies (Coralville, IA).

For probe synthesis, 16 50 µL PCR reactions were conducted in parallel, each with 1 ng oligo pool input using 0.5 µL Phusion polymerase with HF buffer (Thermo Fisher Scientific, Waltham, MA) 1 uM of each primer, and 0.2 mM dNTPs (Thermo Fisher Scientific, Waltham, MA). Cycling conditions were initial denaturation at 98°C for 30s, 12 cycles of 98°C for 10s, 60°C for 30s, and 72°C for 15s, and final extension at 72°C for 10m. Reactions were purified with the QIAQuick Nucleotide Removal Kit (Qiagen, Hilden, Germany), pooling eight reactions per column and eluting each in 30 µL. The elutions were then pooled, and their concentrations were quantified via Qubit 1X dsDNA HS assay (Thermo Fisher Scientific, Waltham, MA).

Four restriction 50µL endonuclease treatments were then performed in parallel, each with 2 µg PCR input and 2 µL FastDigest^®^ LguI (Thermo Fisher Scientific, Waltham, MA). These reactions were then incubated at 37°C for two hours, followed by heat inactivation at 65°C for 5 min. Parallel reactions were then pooled over a single Qiagen MinElute^®^ PCR Purification column (Qiagen, Hilden, Germany), eluted in 10µL, and quantified via Qubit 1X dsDNA HS assay.

Finally, T7 transcription reactions were performed with up to 1 µg of purified LguI digest product, though similar yields were achieved with as little as ∼250 ng. For this HiScribe^®^ T7 High Yield RNA Synthesis Kit (New England Biolabs, Ipswitch, MA) was used according to the manufacturer’s instructions, with one-third of the UTP concentration comprised of Bio-16-UTP (Thermo Fisher Scientific, Waltham, MA). The 20 µL reactions were incubated for 16h at 37°C, after which 68 µL RNase-free H2O was added with 10 µL DNase I Buffer and 2 µL DNase I (New England Biolabs, Ipswitch, MA). The resulting mix was incubated for 15m at 37°C, after which the RNA probes were purified using the Monarch^®^ RNA Cleanup kit (50 µg) (New England Biolabs, Ipswitch, MA). Finally, concentrations were quantified via Nanodrop (Thermo Fisher Scientific, Waltham, MA), and probes were diluted to 100 ng/µL. 100 ng of each probe set was then analyzed on a 12.5% Urea-PAGE gel stained with SYBR-Gold (Thermo Fisher Scientific, Waltham, MA).

### Samples

Two wastewater samples and three soil samples were selected for probe validation with sequencing. The wastewater samples were 24h aggregate influent samples from the city of Hamilton (Ontario, Canada) Wastewater Treatment Plant (November 7, 2022 and March 20, 2023). DNA was extracted from 50 mL wastewater samples within 24h of sampling using the DNeasy^®^ PowerWater Kit (Qiagen, Hilden, Germany) according to the manufacturer’s protocols, each with an associated H2O control. Soil samples were collected from three different environments selected to represent distinct levels of human impact. The first was from Holman Island in the Northwestern Territories of Canada, a pristine environment with little human influence. The second was from a local wetland in an urban setting in Hamilton (Ontario, Canada), representing an environment with middling human impact. The third was from a high-traffic pedestrian area frequented by smokers outside of a Hamilton (Ontario, Canada) hospital, a setting with heavy human influence. DNA was extracted from 250 mg soil samples using the Qiagen DNeasy^®^ Powersoil Kit (Qiagen, Hilden, Germany) according to the manufacturer’s protocols alongside an associated H2O control.

### Sample Processing, Enrichment, and DNA Sequencing

Commercially synthesized probes (CARD v1.0.1 only) and reagents for all enrichments were purchased from Daicel Arbor Biosciences (Ann Arbor, MI). Probes for the allCARD, clinicalCARD, and CARD v1.01 sets were synthesized as outlined above, with the latter allowing direct comparison with commercially synthesized probes. For each sample, extracted DNA was quantified via NanoDrop (Thermo Fisher Scientific, Waltham, MA), diluted, and sonicated to an average size of 400 bp using Covaris G-tubes (Woburn, MA). From this, libraries were prepared in quadruplicate from 1ug DNA using the NEBNext^®^ Ultra II ligation kit (New England Biolabs, Ipswitch, MA) according to the manufacturer’s protocols. These libraries were then pooled before redistributing to half-size indexing reactions using NEBNext^®^ Multiplex Oligos for Illumina (New England

Biolabs, Ipswitch, MA). Sample libraries were then enriched in two batches. The first batched performed allCARD and clinicalCARD enrichments, while the second batch performed commercial and in-house CARD v1.0.1 enrichments (Supplemental Figure 4). All enrichments were performed according to the manufacturer’s protocols (version 5.02) using the Daicel Arbor myBaits v5 kit and reagents in a half-size reaction format with a 24h hybridization at 62°C, maximum library input, and 14 cycles of post-enrichment reamplification. In-house probes were diluted with RNase-free water to the same concentration as Daicel Arbor’s before use in the protocol. Each enriched library was accompanied by an aliquot without enrichment.

Before sequencing, libraries were quantified in triplicate using NEB Luna^®^ Universal Probe qPCR Master Mix (New England Biolabs, Ipswitch, MA), Illumina PhiX standard (San Diego, California) and the following primers, all ordered from Integrated DNA Technologies (Coralville, IA): P5: AATGATACGGCGACCACCGA, P7: CAAGCAGAAGACGGCATACGA, probe: /56-FAM/CCCTACACG/ZEN/ACGCTCTTCCGATCT/3IABkFQ/. Cycling conditions were initial denaturation at 95°C for 3m, followed by 40 cycles of denaturation at 95°C for 15s and annealing/extension at 60°C for 1m. The resulting concentrations were used to make individual pools for each enrichment set above, alongside one replicate of the shotgun (without enrichment) libraries (Supplemental Figure 4). 2x150 bp paired-end reads were obtained from each pool using an Illumina NextSeq 2000 (San Diego, California), with samples having an average depth of 4.47M clusters sequenced (minimum of 3.26M clusters and a maximum of 6.71M clusters).

### Analysis and Visualization

All analyses were performed with custom Python scripts and visualized with the ggplot2 (49) package in R. For rarefaction analysis, libraries were subsampled every 100,000 paired-end reads up to 3M using seqtk v1.3. Fastp v0.23.2 [28] performed initial read trimming, quality control, and deduplication for these subsamples without merging. CARD’s RGI bwt v6.0.2 tool with KMA v1.4.9 (50) then mapped reads to reference sequences in CARD v3.2.5. ARGs were classified as present if they had more than 100 reads mapping (regardless of the breadth of coverage of the reference sequence). Regression lines were determined without extrapolation using the ggplot local estimated scatterplot smoothing function with geom_smooth. Since one cannot extrapolate a locally estimated function, we used a standard log-linear model when extrapolating the predicted number of ARGs detected at a sequencing depth of 10M. GNU parallel v20161222 (51), the Python pandas v1.5.3 library (52), and BioPython v1.78 (53) were used heavily for these analyses.

For read distribution analysis, custom Python scripts used the CARD RGI bwt output to count the number of reads that mapped to each ARG in soil and wastewater samples, respectively. The top 20 most prevalent ARGs (i.e., ARGs with the highest number of reads mapping) in each sample source (soil, wastewater) were kept for the figure, while all others were collapsed to the ‘other’ category. This cut-off was chosen to keep figure legends readable while providing discriminatory power between the performance of different probe sets. Mean counts of the two replicates are used to determine the number plotted. The distribution of percent identity between read and reference for each sample was determined by a custom Python script that parsed the CIGAR strings in the BAM file that accompanies the RGI bwt output, where the percent identity was calculated as the number of nucleotide matches / 151 * 100. Replicates were pooled for this analysis.

In a clinicalCARD versus allCARD overlap analysis, a custom Python script determined clinically relevant ARGs detected by allCARD or clinicalCARD with >=100 mapped reads in both replicates at each subsampling depth. A similar approach was used to determine the overlap between the in-house vs commercially synthesized CARD v1.01 probe sets. However, to compare these sets, only ARGs included in the initial design (i.e., in CARD v1.0.1) were considered. Coverage analysis only considered clinically relevant ARGs detected by both clinicalCARD and allCARD with at least one read at a subsampled sequencing depth of 3M paired-end reads in both replicates. Python scripts determined each ARG’s coverage using the CARD RGI bwt output from the rarefaction analysis. Once again, for a similar analysis comparing the commercial and in-house synthesis, only ARGs from CARD v1.0.1 were considered.

For a wastewater read correlation analysis, only clinically relevant (i.e., in clinicalCARD) ARGs detected with at least one read in both shotgun and enriched in each sample were considered. All ARGs were included from soil samples, while clinicalCARD analysis was omitted due to the sparsity of detected ARGs. R^2^ values were calculated using scikit-learn v1.2.2 (54). The cmlA1 read number correction outlined below was accomplished by summing the number of reads attributed to all closely related cmlA variants (i.e., cmlA1, 4, 5, 6, 8, 9) in each relevant treatment and manually adding the corresponding values to the plot.

## Results

### Probe Design and Synthesis

After design and filtering, allCARD contained 34,915 unique probe sequences covering 4,661 ARGs. Alternatively, clinicalCARD contained 15,393 unique probes, covering 323 ARGs (Supplemental Figure 5). No probe set had zero coverage of an ARG against which it was designed. When analyzing probes against all ARGs curated as CARD protein homolog models (Figure 2A), the median value of all metrics other than the proportion covered per ARG was higher in the clinicalCARD probe set. The reason for clinicalCARD’s bimodal distribution compared to allCARD is that it was only designed against the clinically relevant subset. Therefore, the clinically irrelevant genes (e.g., TolC, H-NS) had no probes aligning since they were not included in the initial design. However, despite only being designed against 323 ARGs, over 50% of CARD ARGs had coverage of >75% by clinicalCARD. When analyzing both probe sets against the clinically relevant ARGs in clinicalCARD (Figure 2B), median values in clinicalCARD were higher than allCARD in every metric.

**Figure 2:**
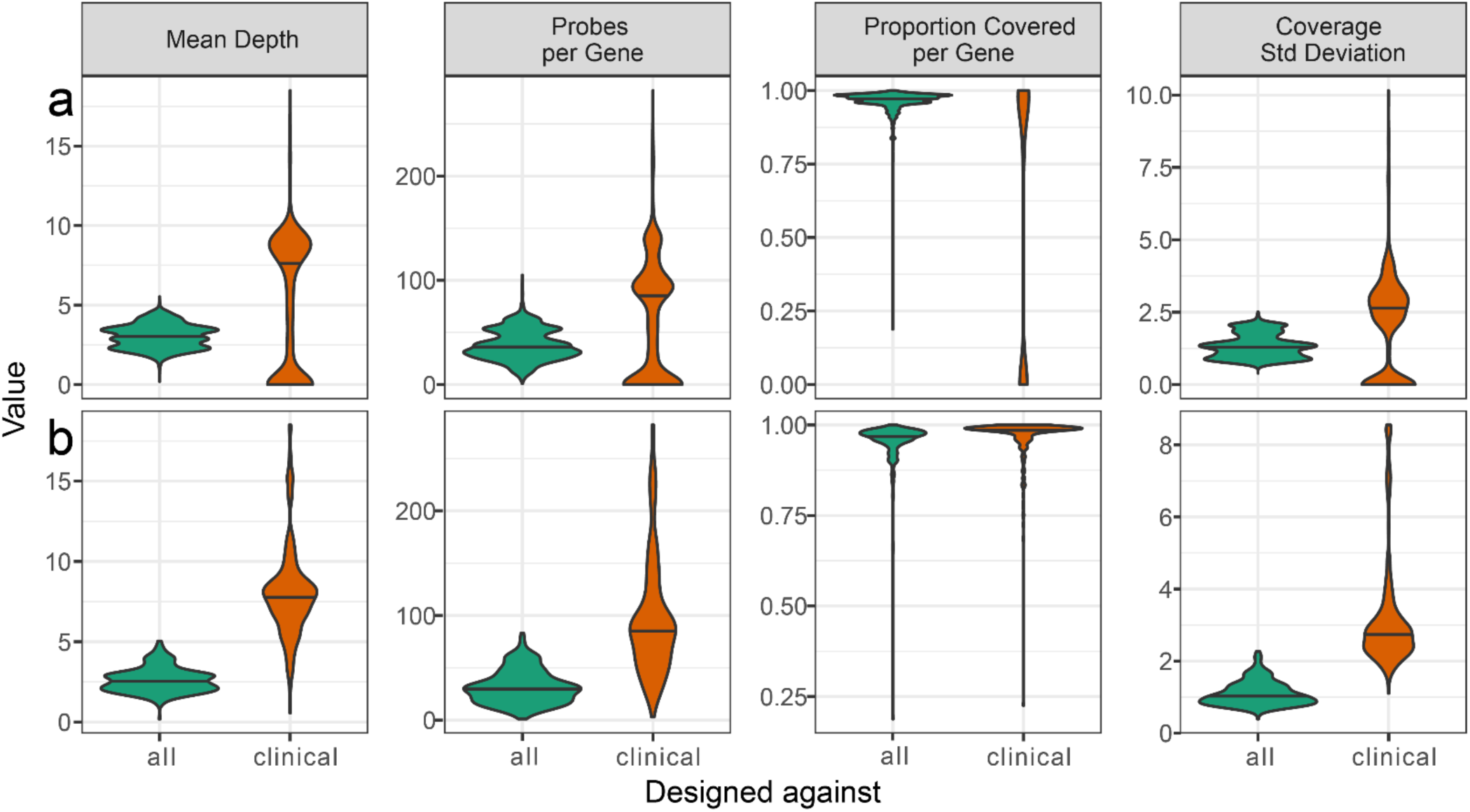
*In-silico* probe analysis using BLAST to align all probes in allCARD and clinicalCARD against a) all genes in CARD and b) all genes deemed to be clinically relevant.

The increased *in silico* performance of clinicalCARD is a consequence of the smaller initial set of reference sequences. As such, it did not have to remove as many probes during the redundancy filter (Supplemental Figure 5A) and ended up with more redundancy and superior coverage of the ARGs against which it was designed. Probes in clinicalCARD had a lower median GC content and melting temperature. A higher proportion of probes also target only a single ARG in clinicalCARD relative to allCARD (Supplemental Figure 5B). After synthesis, a urea-PAGE gel shows a smear above 80 nt due to the stochastic incorporation of biotin into the probe (Supplemental Figure 6). A comparison of all pools against the commercially synthesized version indicates a similar size distribution, with a slight bias towards lower molecular weights in the in-house synthesized probe set.

### Probe Validation

Five samples in five different treatments were employed to determine the efficacy of the probes when enriching for targets. No blanks had sufficient sequencing depth to be analyzed at even the lowest subsampling depth, indicating negligible contamination. First, the non-inferiority of the in-house synthesized probe set relative to the commercial option was established. After subsampling every 100k paired-end reads up to 3M with analysis by the CARD RGI bwt tool (55), the number of ARGs with >100 mapped reads was determined for each depth in each sample (Figure 3). This analysis illustrated that the in-house synthesized probes detected more ARGs at the same sequencing depth in every sample. This effect was more pronounced in the soil samples, where the in-house synthesized probe set detected as many as double the number of ARGs detected by the commercially synthesized probe set, despite both having identical probe sequences based on CARD v1.01. Yet, our in-house synthesized probe set had an almost identical distribution of the top 20 detected ARGs after enrichment compared to the commercially synthesized set (Figure 4), indicating consistent enrichment efficiency between ARGs in the commercial and in-house synthesized sets. These results were further complemented by analyzing the overlap between ARGs detected by each set at each subsampling depth and the associated coverage distribution for all detected ARGs (Supplemental Figures 7 & 8). In these analyses, enrichment with our in-house synthesized probes detected ARGs with less sequencing effort than commercially synthesized probes and had greater coverage of detected ARGs.

**Figure 3:**
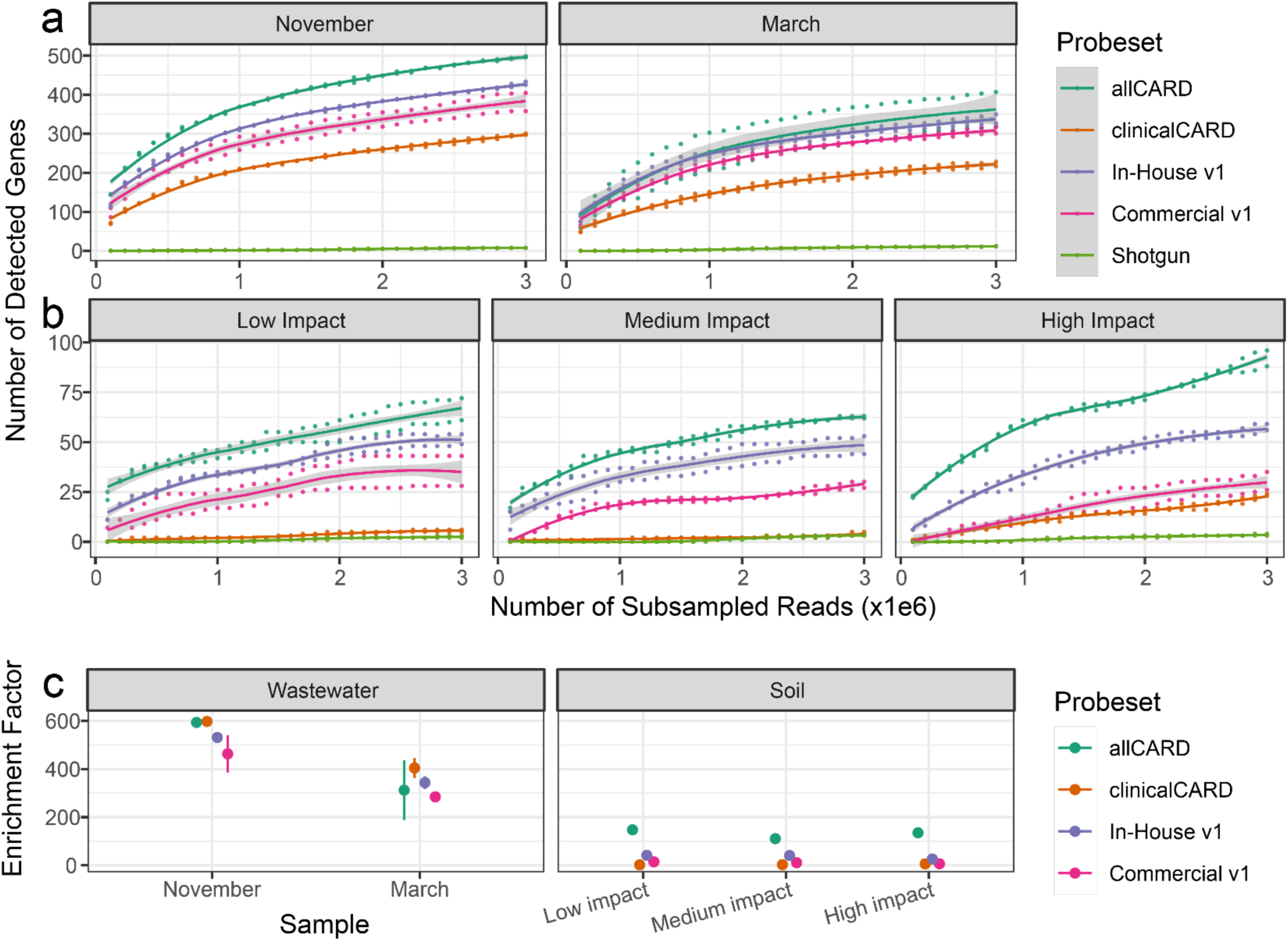
Enrichment efficiency comparisons between different treatments. a) Rarefaction analysis of the number of detected genes at subsampling depths of every 100k reads up to 3M in wastewater samples and b) soil samples. Shaded areas represent the 95% confidence interval. c) Average enrichment factor ±1SD of different probe sets in different samples. The enrichment factor is defined here as the number of CARD-mapped reads after enrichment with the relevant probe set divided number of CARD-mapped reads in shotgun sequencing data.

**Figure 4:**
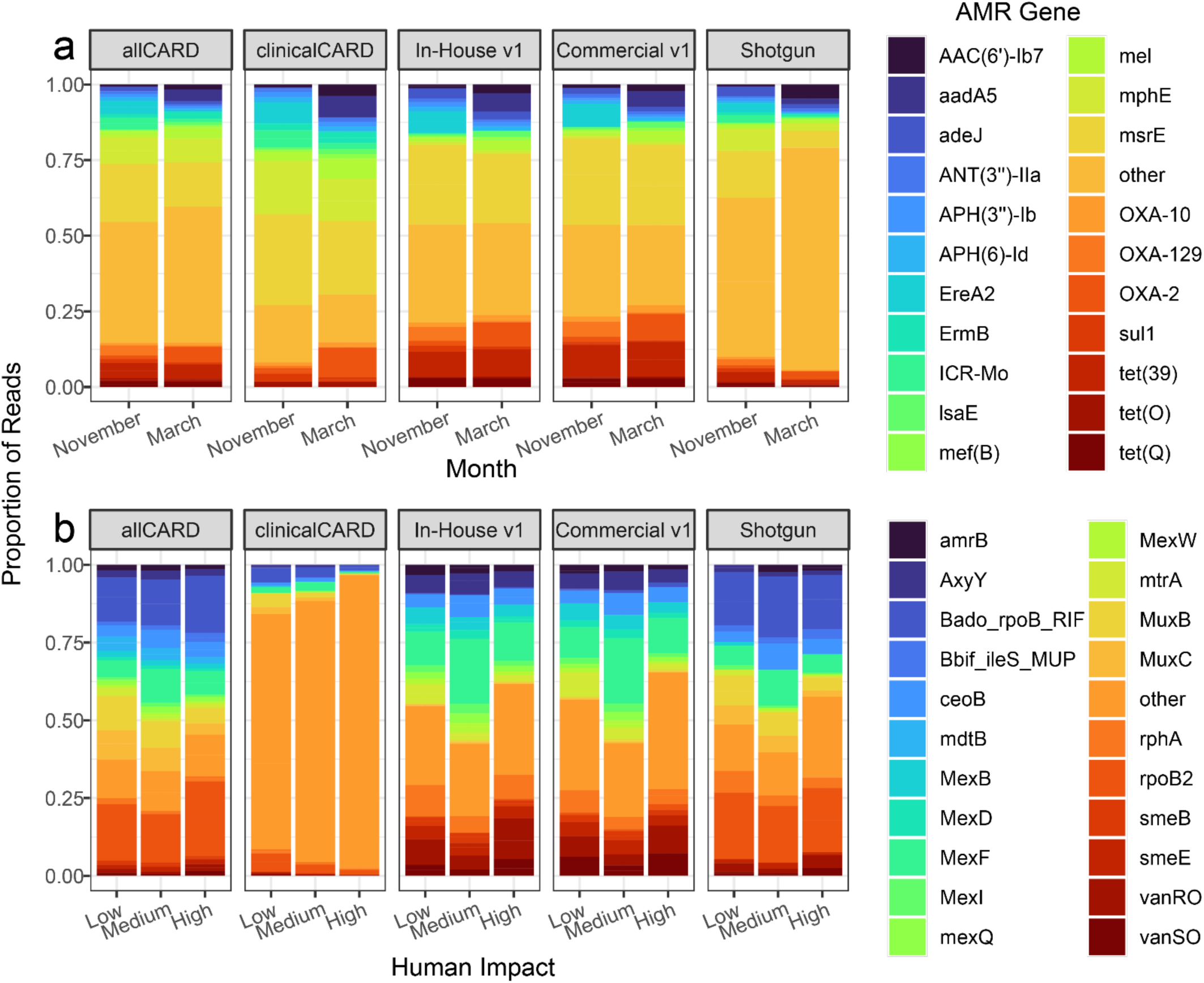
Top 20 AMR genes according to overall count and their associated read proportions in each sample. a) Wastewater samples. b) Soil samples. Very long gene names were replaced with their CARD short name.

Enrichment with all probe sets detected vastly more ARGs in wastewater samples than by sequencing without enrichment. AllCARD detected the most, with up to 498 different ARGs, and clinicalCARD detected the least, with up to 300 ARGs. However, clinicalCARD efficiently retained the highest abundance ARGs, evidenced by the similar distributions between allCARD and clinicalCARD in Figure 4A. AllCARD detected by far the greatest number of ARGs in soil samples, up to 96 in the high-impact hospital grounds. ClinicalCARD did not detect a substantial number of ARGs in any soil sample, except for that taken from the high-impact hospital grounds, where it detected 24.

The enrichment factor (i.e., the number of enriched reads mapped to CARD divided by the number of shotgun reads mapped to CARD) of different probe sets shows a more consistent value within each sample than expected, given the high performance of allCARD at detecting the most ARGs (Figure 3C). This is especially evident in wastewater, where clinicalCARD is on par with, if not outperforming, allCARD in terms of enrichment factor despite detecting far fewer ARGs in the rarefaction curve. There was a 400 to 600-fold enrichment in the wastewater samples in the November sample, depending on the probe set, which decreased to 200 to 400-fold for the March sample. Soil samples had a markedly lower enrichment factor than wastewater, hovering between 0 to 200-fold enrichment depending on the probe set. Yet, enrichment was consistent between samples for different probe sets in soil. Based on the rarefaction curves (Figure 3), ARG detection in most samples begins to plateau by a sequencing depth of 3M paired-end reads. Extrapolation of these rarefaction curves to a sequencing depth of 10M paired-end reads indicates that at a subsampling depth of 3M, we have captured 70%-80% of the diversity of ARGs that may be detected at 10M (Supplemental Figure 9).

There were differences when comparing the top 20 most prevalent ARGs in soil and wastewater (Figure 4). First, all top 20 ARGs in wastewater data, save for adeJ and tetQ, were in our set of clinically relevant ARGs. However, in soil samples, not a single top 20 ARG was included in the clinically relevant set, which explains the lack of enrichment for any top 20 ARGs in soil by the clinicalCARD probe set. Most dominant ARGs in the soil samples were found across various species and associated with efflux pumps (e.g., Mex proteins) or transcriptional regulators (e.g., mtrA, vanR/S). Moreover, the percent identity of the ARG-associated reads relative to the CARD reference sequences in the soil samples was lower than those from wastewater samples (Figure 5). Before enrichment, the wastewater sample had a mixture of high-identity (>90%) and mid-identity (60%-90%) reads relative to their references in CARD, but the soil samples had exclusively mid-identity to low-identity (<60%) reads. After enrichment, wastewater samples had almost exclusively high-identity reads in both the clinicalCARD and allCARD enrichments. In soil samples, enrichment with allCARD selected heavily for mid-identity reads, but clinicalCARD preferentially selected for high-identity reads, especially in the sample from the high-impact hospital grounds.

**Figure 5:**
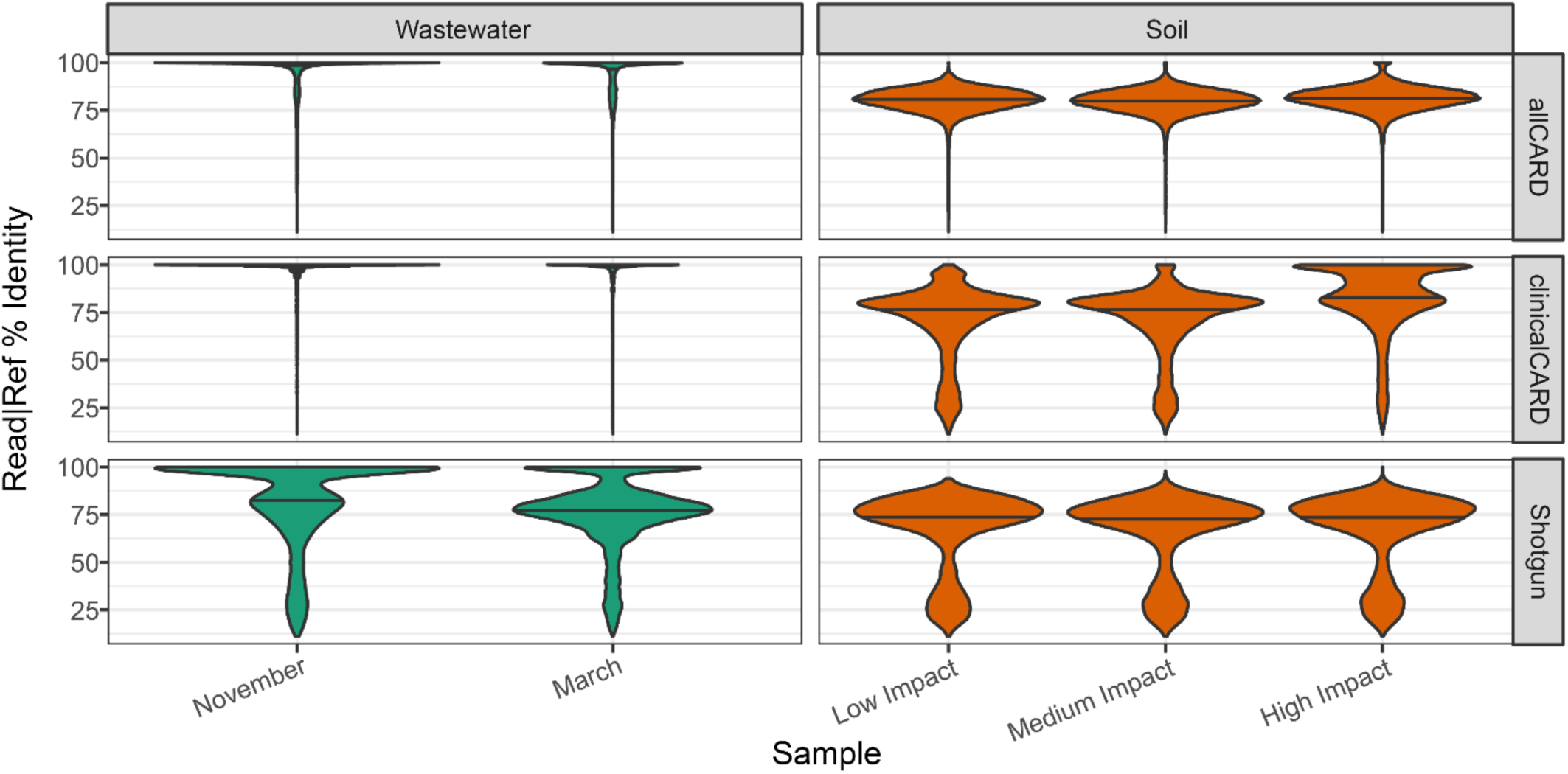
Distribution of the number of similarity between read and reference, based on CIGAR strings from the BAM file output by the CARD RGI bwt module. The internal line of each violin plot indicates the median of the distribution.

### AllCARD versus ClinicalCARD

The wastewater samples were used to examine clinicalCARD’s efficacy relative to allCARD at enriching the clinically relevant ARGs it was designed against, as clinically relevant ARGs had the highest abundance in the wastewater samples. Overall, clinicalCARD detected an average of 16% more ARGs than allCARD in the November wastewater sample and 35% more in the March sample at each subsampling depth, with very rarely an ARG detected uniquely by allCARD (Figure 6A).

**Figure 6:**
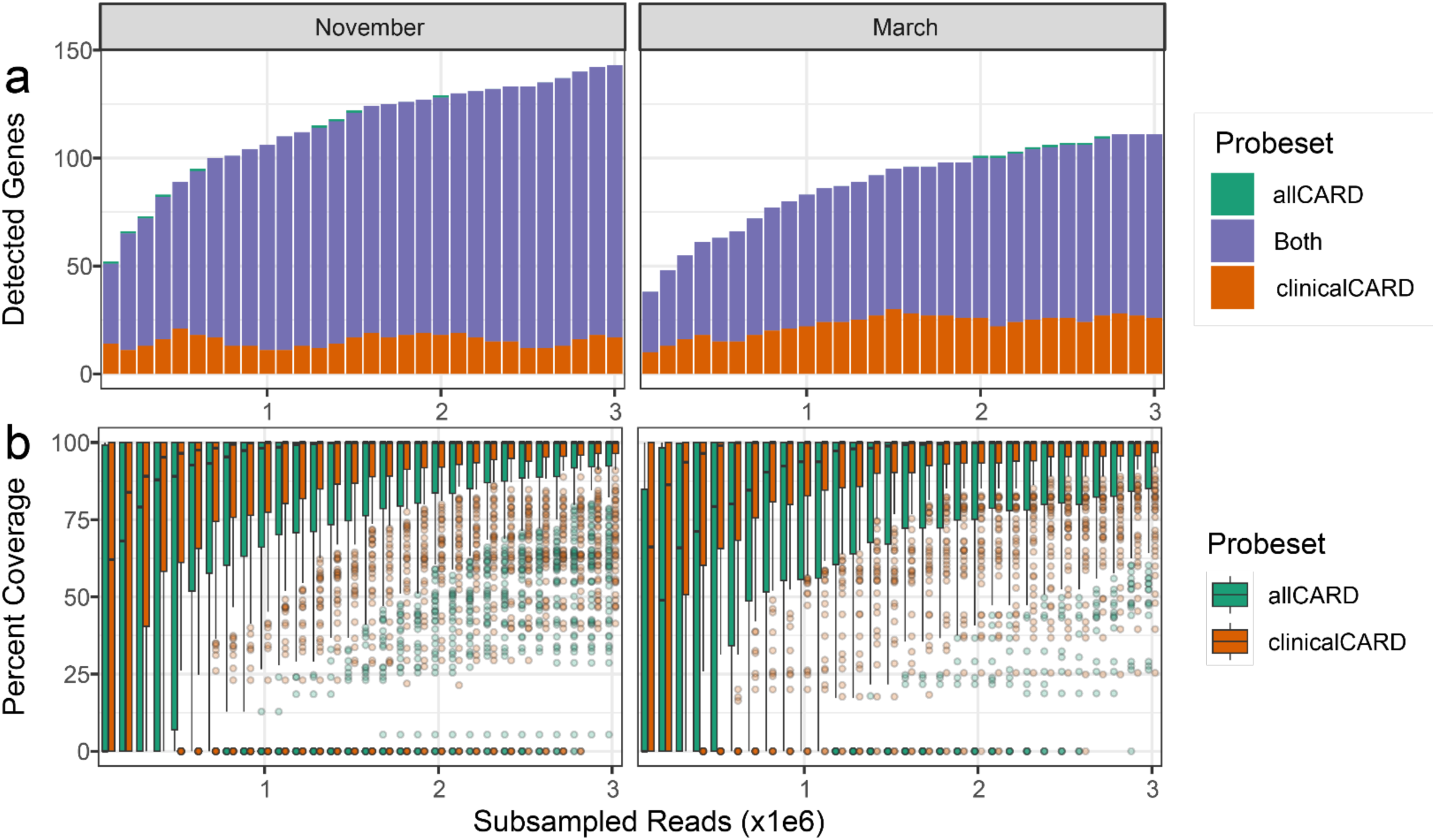
Comparison of enrichment efficiency of clinically relevant ARGs between allCARD and clinicalCARD probe sets. **a)** Overlap analysis showing ARGs detected by allCARD, clinicalCARD, for both with at least 100 reads at different subsampling depths. **b)** Coverage analysis showing the distribution of coverage for ARGs detected by both probe sets with at least one read at a depth of 3M. In the November sample, there were 226 such ARGS. In the March sample, there were 176.

To assess the breadth of coverage of individual ARGs after enrichment with allCARD or clinicalCARD, clinically relevant ARGs detected by both probe sets in both replicates at a subsampling depth of 3M paired-end reads were analyzed. The distribution of coverage of these ARGs at each subsampling depth for allCARD and clinicalCARD was plotted (Figure 6B). ClinicalCARD delivered 100% coverage of at least 50% of the ARGs detected in the November sample at a subsampling depth of 1M reads, while allCARD took 1.5M reads to do the same. In the March sample, clinicalCARD took only 700k reads to reach this level of detection, while allCARD required 2.1M reads. Based on this analysis, clinicalCARD delivers better coverage of more clinically relevant ARGs at a lower sequencing depth than allCARD.

Finally, a commonly perceived limitation of enrichment is that hybridization can be sequence-dependant, which may introduce biases in the final library, thereby eliminating the ability to perform relative quantification of ARGs in a sample. To investigate the relationship between ARG abundance in enriched versus unenriched data, the clinically relevant ARGs present with at least one read in both replicates of shotgun and enriched data at a subsampling depth of 3M paired-end reads were analyzed (Figure 7). In the wastewater samples, due to the high identity between reads, the R^2^ value reached as high as 0.996 in the November sample with the allCARD probe set, while it was slightly lower after enrichment with clinicalCARD. In the March wastewater sample, R^2^ values were considerably lower, most likely due to the inconsistent replicates. Enrichment was generally less efficient in the soil samples due to the lower identity between DNA and probes; however, the R^2^ values remained high, reaching 0.897 in the low-impact sample, 0.936 in the medium-impact, and 0.775 in the high-impact sample from the hospital grounds.

**Figure 7:**
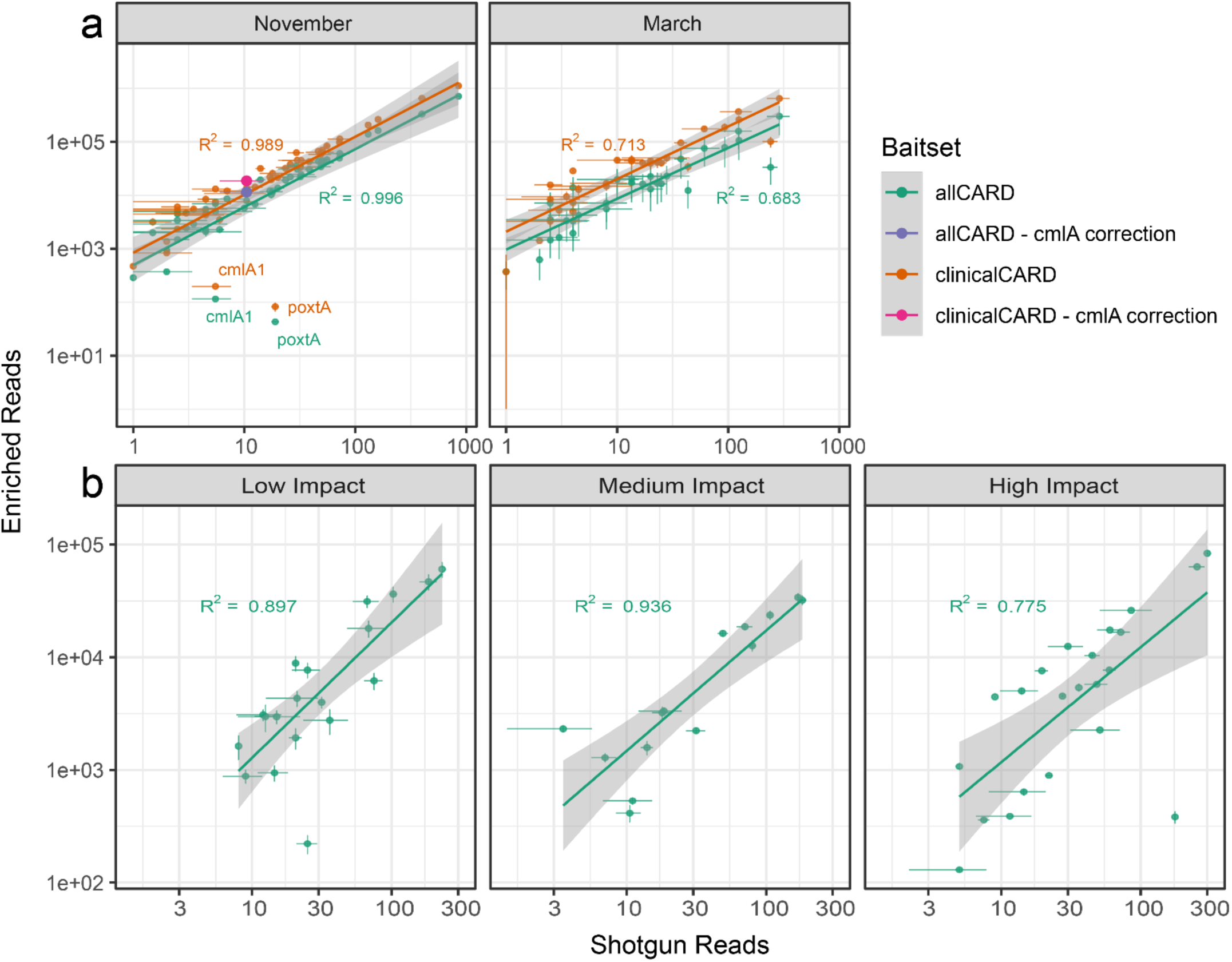
Correlation of the number of enriched and shotgun reads for each ARG with associated R^2^ values. Shaded areas represent the 95% confidence interval. **a)** Wastewater analysis: 44 and 31 ARGs fit inclusion criteria for the November and March samples, respectively. The purple and pink dots show cmlA1’s position for allCARD and clinicalCARD enrichment, respectively, if all reads from all extremely closely related cmlA variants were considered to be from cmlA1. **b)** Soil analysis: clinicalCARD was not analyzed due to a lack of reads mapping to clinically relevant ARGs. In the low human impact sample, 19, 14, and 22 ARGS fit the inclusion criteria for the low human impact, medium impact, and high impact samples, respectively.

In Figure 7, ARGs were labelled on the plot if the enrichment factor was significantly lower (p < 0.05) than the average enrichment factor within a sample and probe set. Two ARGs in the November wastewater sample were obvious outliers. The first, cmlA1, has several close homologs with >99% nucleotide identity to which thousands of reads were assigned. This occasional occurrence of reads mapping among highly similar alleles is known as the allele network problem (50, 56). When mapping to highly redundant databases, even with new tools such as KMA (50) as used by RGI bwt, reads can be misassigned to another closely related allele or ARG. To illustrate this, we superimposed corrected points onto the plot, i.e., showing where cmlA1 would reside on the plot if all the reads attributed to its variants were attributed to it instead. As expected, this correction brings it into far better agreement with the observed trend. The reads of the second outlier ARG, paxtA, appear to be from a distant homolog of the reference in CARD. Due to this sequence variance, it has a lower enrichment efficiency.

## Discussion

Our work has resulted in the design and validation of two probe sets to enrich ARGs in metagenomic sequencing libraries. Several improvements have been made relative to the first iteration of this probe set against the first version of CARD. First, a denser initial tiling and exclusion of perfect matches during the nt BLAST filter maximized coverage against all ARGs. Second, a stringent redundancy filter minimized unneeded probes. This enables allCARD to enrich all ARGs included in CARD more evenly with fewer probes. Alternatively, clinicalCARD has a much higher redundancy since it was designed against a much smaller set of clinically relevant ARGs. Our in-house synthesis protocol yields a similar smear pattern to the commercially synthesized version when run on a urea-PAGE Gel, indicating similar physical properties. There was a slight bias to lower molecular weights in the in-house synthesized probe sets, which may indicate more incorporation of biotin into the commercial probes.

When comparing *in silico* statistics of allCARD versus clinicalCARD, it is evident that clinicalCARD outperforms allCARD when enriching for the ARGs it was designed against. ClinicalCARD is superior at targeting the ARGs against which it was designed due to its decreased need for redundancy filtering during probe design. Since it covers a smaller number of ARGs, fewer probes are made during the initial tiling step with BaitsTools, and fewer cycles of redundancy filtering are needed to satisfy the total probe number cut-off (Figure 2A). This translates to more probes per ARG and, therefore, better coverage of the ARGs against which it was designed. However, clinicalCARD also has coverage against a much larger portion of CARD than only those ARGs (i.e., median coverage of clinicalCARD probes against allCARD ARGs is >75%). This is because of the high degree of conservation of ARG nucleotide sequences and the resulting redundancy in CARD, where one ARG may have several closely related variants (35). Overall, ClinicalCARD has higher median scores in all tested metrics against its design set of ARGs. Based on this, clinicalCARD should be more effective when enriching clinically relevant ARGs. Additionally, since it is a smaller set of probes, it will be less expensive, even with in-house synthesis. However, it lacks coverage of many ARGs in allCARD. As some questions require comprehensive coverage of all ARGs, allCARD is the better choice in these situations.

To test our new probe sets, five samples were used for validation: two wastewater samples spanning the beginning and end of winter and three soil samples from differently human-impacted sites. In all samples, enrichment drastically improved the detection of ARGs relative to shotgun sequencing, with an enrichment factor of up to 598-fold. Our in-house synthesis consistently detected more ARGs at a lower sequencing depth than the commercially synthesized probes. This may be due to the evenness of coverage in the Twist Biosciences oligo-pool, which may provide a superior template for transcription and, thus, a superior capture reagent.

In wastewater samples, allCARD detected the most ARGs by a considerable margin. ClinicalCARD detected relatively few ARGs, though those it did detect were among the most frequent ARGs in the sequencing data. AllCARD once again detected the most ARGs in the soil samples, but clinicalCARD missed many ARGs in soil except in the sample from the high-impact hospital grounds. This is likely due to the lack of clinically relevant ARGs in soil samples from environments with less human impact. We stress, however, that this experiment, as designed, does not indicate causality from being near a hospital. It merely supports the notion that samples from environments with high human influence carry more ARGs, a known phenomenon (57, 58).

Compared to the large fluctuation in the number of ARGs detected between probe sets, the enrichment factor among probe sets was more consistent, especially in the wastewater samples. This suggests that with a certain input of probes, one can expect a given level of enrichment so long as the target is sufficiently abundant. Enrichment suffers when this is not the case, such as in the non-allCARD probe sets for the soil samples. When inspecting the number of identities between reads and references, there is a pronounced difference between wastewater and soil samples. Without enrichment, the distribution of the number of identities in unenriched reads of wastewater samples indicates the presence of both distant and close homologs to the references in CARD. After enrichment, only close homologs are present, suggesting that they are preferentially enriched. However, distant homologs are enriched in their absence, such as in the low- and medium-impact soil samples (Figure 5B). However, in the sample from the high-impact hospital grounds, there was a bias towards close homologs, especially in the sample enriched by clinicalCARD. Overall, this shows a pattern where clinically relevant ARGs have a higher identity to their references in CARD and thus are more efficiently enriched, particularly given that CARD and other ARG databases are biased towards clinical isolates (35). The distant homologs in soil samples are mainly associated with efflux and regulation, and we cannot know that these distant homologs confer the same degree and type of AMR as their reference in CARD. For this, experimental data are required.

There was a striking linearity when considering the correlation between the number of ARG-associated reads in enriched versus unenriched samples. This was unexpected due to the assumed effect that sequence differences, specifically GC content, would have on hybridization efficiency (59, 60). Relative quantification of ARGs within a sample and comparison among samples may be possible, i.e., a large difference in read abundance in enriched data indicates a proportionally large difference in shotgun data, although smaller differences may be challenging to detect reliably. However, our investigation contains few samples, and quantifying this relationship was not our primary aim.

There are limitations to enrichment as a method for resistome profiling. Enrichment cannot detect entirely novel resistance genes that are not in CARD. As such, we designed CARPDM to update the probe set for each released version of CARD. Combined with reduced up-front costs via our novel synthesis method, researchers can update their probe set on demand. Moreover, when the genes that get enriched are distant homologs to those in CARD, we cannot be sure if they are actual resistance genes or if they would generate clinical levels of resistance. More work is required to validate these genes upon detection. Finally, this protocol relies on proprietary hybridization reagents and buffers from commercial suppliers.

## Conclusions

The increasing global burden of AMR requires cost-effective and scalable solutions. Overall, our in-house synthesized probes detect more ARGs with less sequencing than a commercial option in the samples tested. AllCARD robustly enriches the vast array of ARGs against which it was designed, as well as their distant homologs. ClinicalCARD even more robustly enriches the smaller set of clinically relevant ARGs against which it was designed. This work shows that targeted enrichment is a valuable companion to DNA sequencing when detecting ARGs. Future work will involve further updating and refining the set of clinically relevant ARGs and continually updating the probe set with every new CARD release. Additionally, while we have released a preliminary protocol for the in-house synthesis of any probe set, there remains room for this protocol to be optimized. Finally, to make this technology more accessible, future work should also investigate its efficacy on alternative DNA sequencing platforms, such as Oxford Nanopore’s MinION. Yet, overall, this work has shown the power of enrichment to decrease the cost and increase the impact of large-scale monitoring of ARGs using DNA sequencing. Alongside CARD, this technology can help researchers investigate ARG prevalence and transmission patterns among different populations and environments.

## Data, Protocol, and Probe Sequence Availability

Detailed laboratory protocols and FASTA sequence files for probe synthesis and library enrichment are available at the Comprehensive Antibiotic Resistance Database website: http://card.mcmaster.ca.

## Acknowledgments

Computational support was provided by the McMaster Service Lab and Repository (MSLR) computing cluster, supplemented by hardware donations and loans from Cisco Systems Canada, Hewlett Packard Enterprise, and Pure Storage. AGM holds McMaster’s inaugural David Braley Chair in Computational Biology, generously supported by the family of the late Mr. David Braley. Additional thanks go to the McMaster Genomics Facility for next-generation sequencing and all the members of the McMaster Ancient DNA Center.

