## Supplementary Figures 1-9 for "CARPDM: cost-effective antibiotic resistome profiling of metagenomic samples using targeted enrichment"

### Supplemental Figures

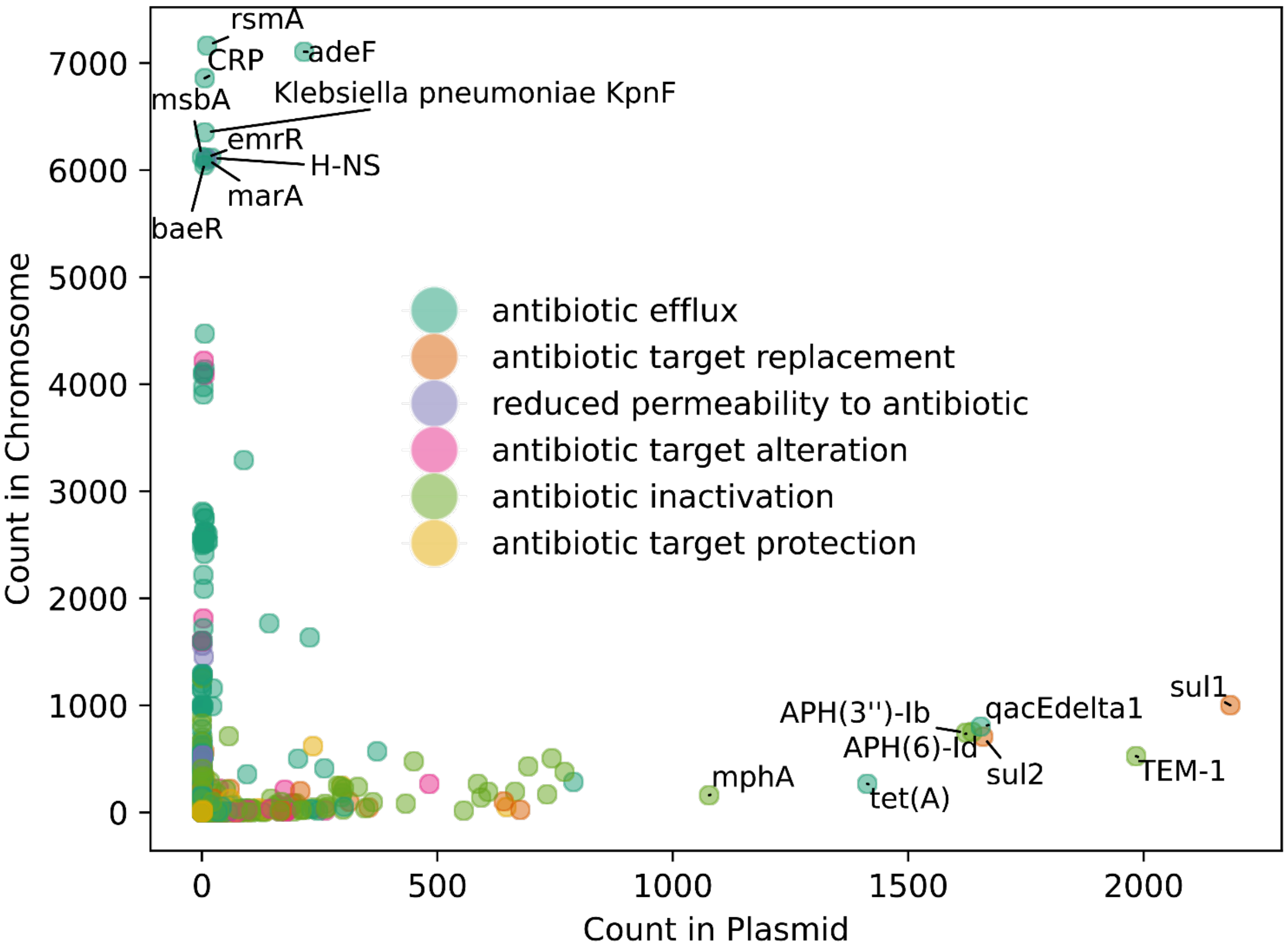


Supplemental Figure 1: Prevalence data for plasmids and chromosomes for ARGs in CARD. ARGs are coloured according to their annotated AMR mechanism in CARD.

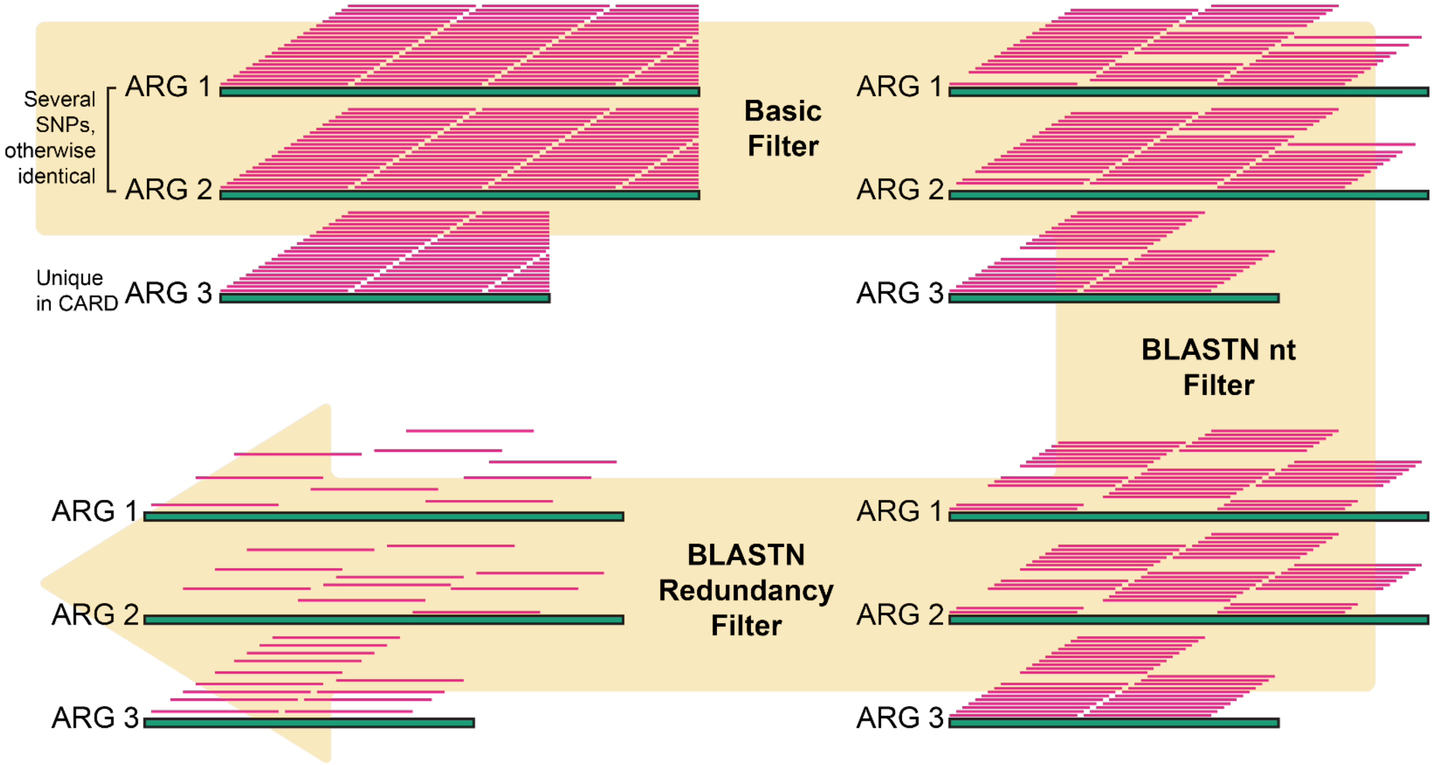


Supplemental Figure 2: CARPDM probe design strategy. The workflow starts with 80 nt long probes tiled every 4 nt along all sequences in the input, constructed by Baitstools (44). In this simplified example, there are two highly similar ARGs and one that is unique in CARD, with no close homologs. During the basic filter, probes are filtered out if they don't meet specific physical characteristics (e.g. length of 80 bp, melting temperature > 50°C, no ambiguous bases, etc.). Probes are also deduplicated, as redundant sequences in the design aren't helpful during enrichment. Next, the probes are removed based on if they have hits in the BLASTN nt database. Finally, redundant probes are iteratively collapsed based on their sequence similarity until the total number of probes is below a defined cut-off. Note that ARGs 1 and 2 have lower probe density than ARG 3. However, due to 1 and 2's similarity, each probe designed against one will also hybridize with the other, leading to a similar final probe density as 3.

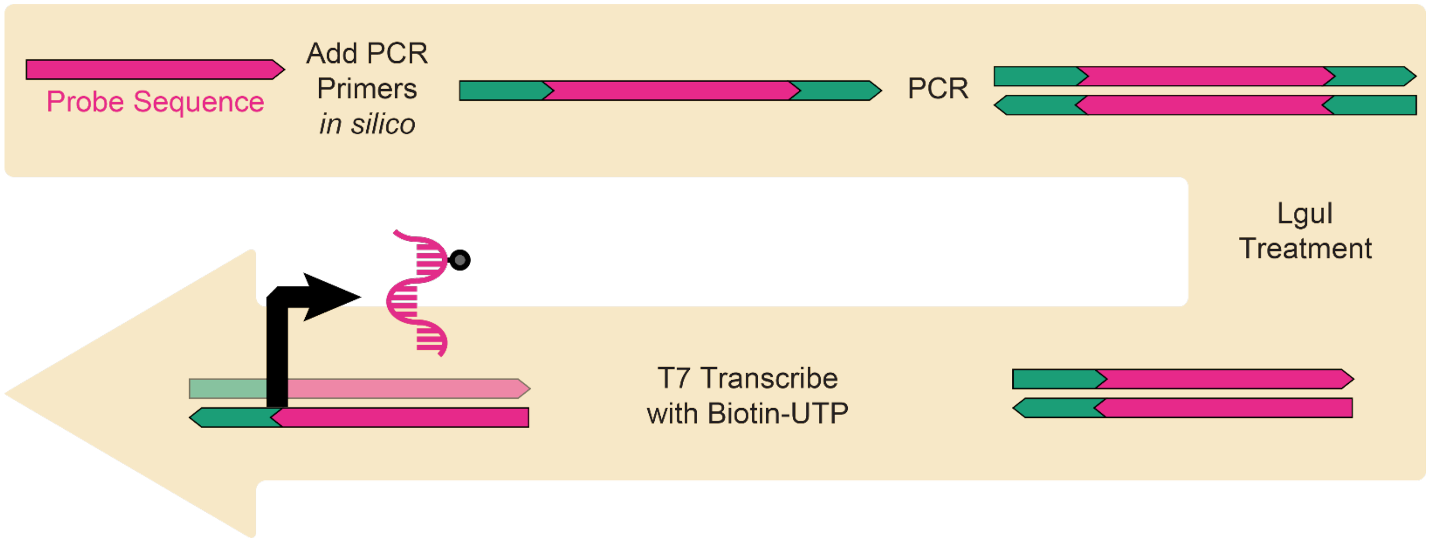


Supplemental Figure 3: Probe synthesis workflow. The first step is to add amplification primers to either end of every unique probe sequence in the set. This allows PCR amplification of the synthesized oligo pool before LguI restriction endonuclease treatment and subsequent T7 RNA transcription in the presence of biotinylated UTP.

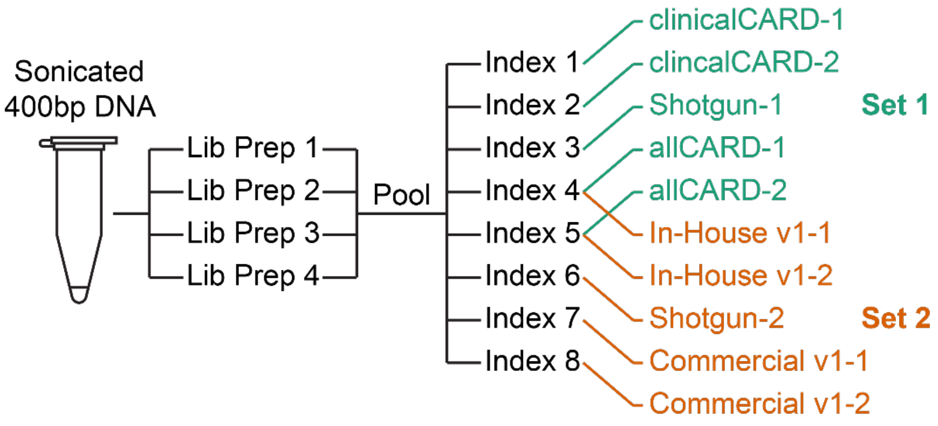


Supplemental Figure 4: Experimental design, indicating the workflow and separation of library preparation, indexing, and enrichment treatments. Each set was sequenced on its own sequencing run to gain sufficient depth.

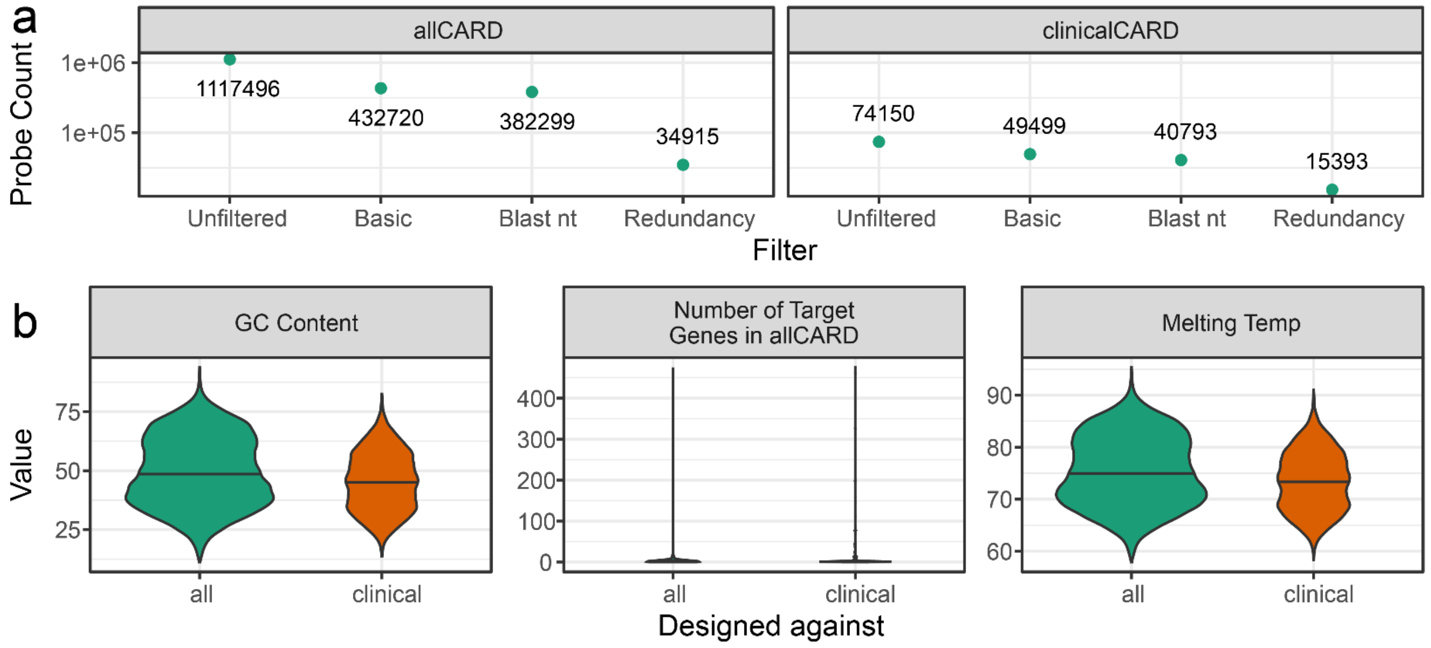


Supplemental Figure 5: AllCARD and clinicalCARD probe set statistics. a) Number of probes remaining after each filter employed during design. b) Physical statistics of probes in each set.

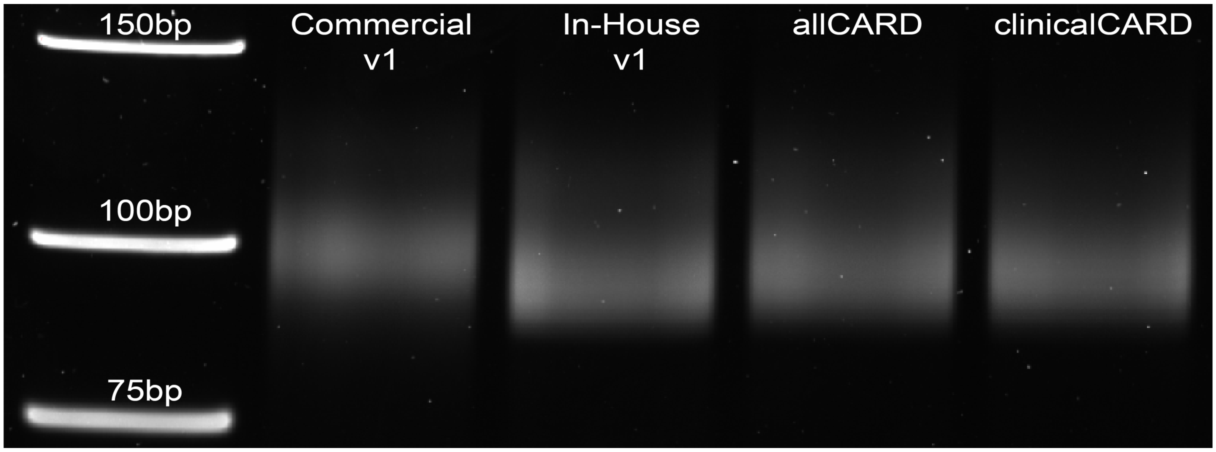


Supplemental Figure 6: 12.5% Urea-PAGE gel comparing size distributions of commercial and in-house synthesized probe sets.

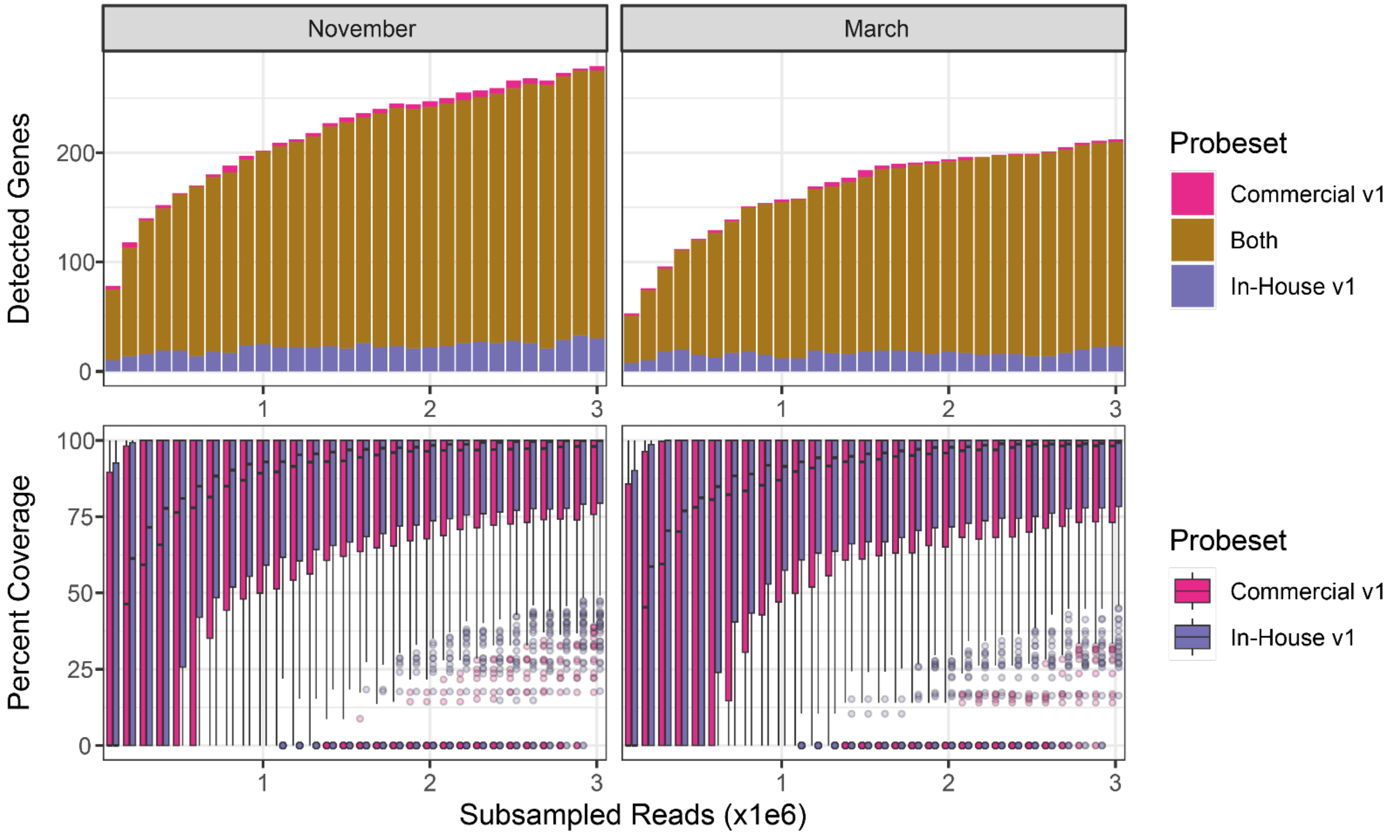


Supplemental Figure 7: Analysis of overlap and ARG coverage distribution in wastewater samples after enrichment with the commercial and in-house synthesized CARD v1.0.1 probe sets. ARGs considered are limited to those included in CARD v1.0.1, used for the initial design of this probe set.

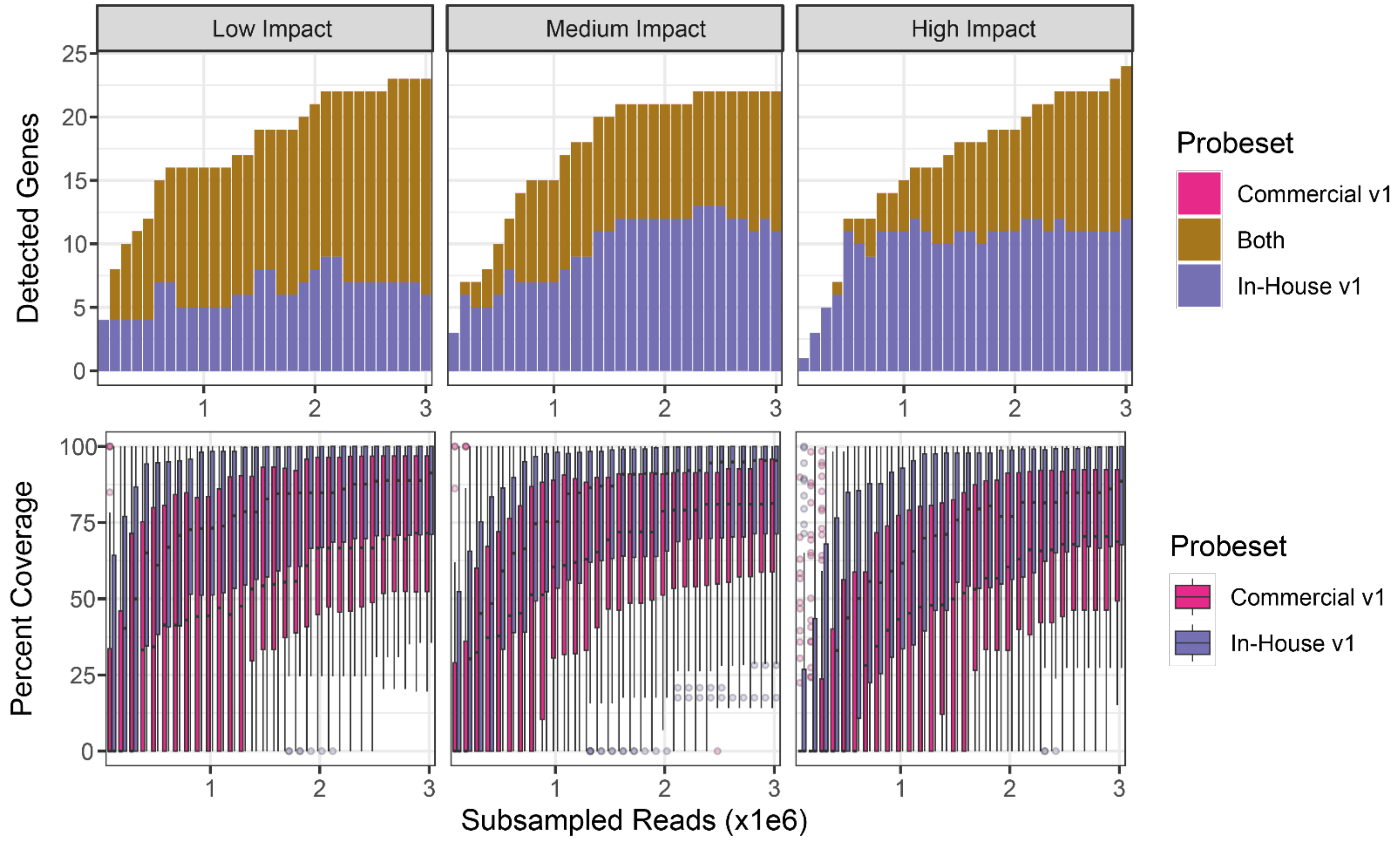


Supplemental Figure 8: Analysis of overlap and ARG coverage distribution in soil samples after enrichment with the commercial and in-house synthesized CARD v1.01 probe sets. ARGs considered are limited to those included in CARD v1.0.1, used for the initial design of this probe set.

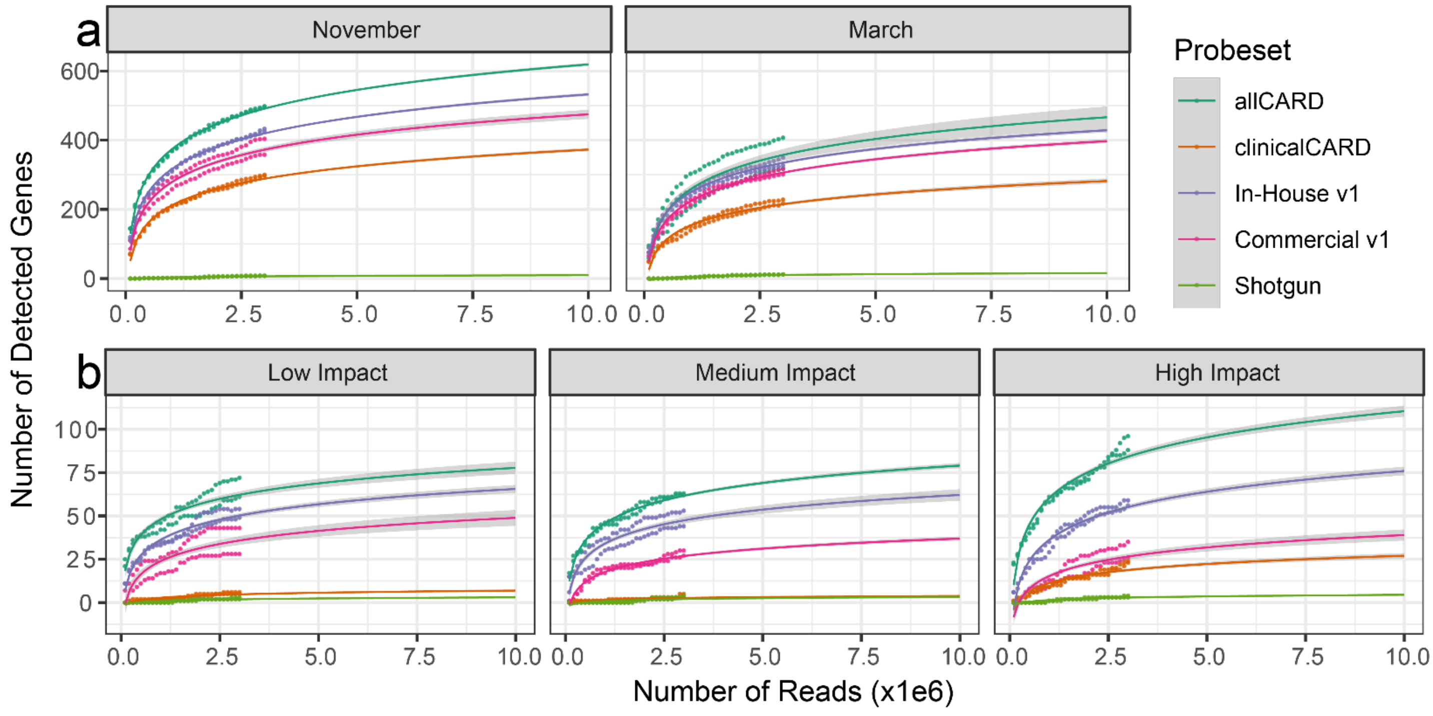


Supplemental Figure 9: Rarefaction curves from Figure 3 extrapolated to show the number of detected ARGs at 10M reads. Shaded areas represent the 95% confidence interval.
